# Identifying, understanding, and correcting technical biases on the sex chromosomes in next-generation sequencing data

**DOI:** 10.1101/346940

**Authors:** Timothy H. Webster, Madeline Couse, Bruno M. Grande, Eric Karlins, Tanya N. Phung, Phillip A. Richmond, Whitney Whitford, Melissa A. Wilson Sayres

## Abstract

Mammalian X and Y chromosomes share a common evolutionary origin and retain regions of high sequence similarity. This sequence homology can cause the mismapping of short sequencing reads derived from the sex chromosomes and affect variant calling and other downstream analyses. Understanding and correcting this problem is critical for medical genomics and population genomic inference. Here, we characterize how sequence homology can affect analyses on the sex chromosomes and present XYalign, a new tool that: (1) facilitates the inference of sex chromosome complement from next-generation sequencing data; (2) corrects erroneous read mapping on the sex chromosomes; and (3) tabulates and visualizes important metrics for quality control such as mapping quality, sequencing depth, and allele balance. We show how these metrics can be used to identify XX and XY individuals across diverse sequencing experiments, including low and high coverage whole genome sequencing, and exome sequencing. We also show that XYalign corrects mismapped reads on the sex chromosomes, resulting in more accurate variant calling. Finally, we discuss how the flexibility of the XYalign framework can be leveraged for other use cases including the identification of aneuploidy on the autosomes. XYalign is available open source under the GNU General Public License (version 3).

## Introduction

Accurate genotyping and variant calling are priorities in medical genetics, including molecular diagnostics, and population genomics (Taylor *et al*., 2015; Ashley, 2016). Despite the availability of numerous powerful tools developed to infer genotypes from sequencing data, sequence homology among genomic regions still presents a major challenge to genome assembly, short read mapping, and variant calling. Specifically, similar sequence content can confound the mapping of short next-generation sequencing reads to a reference genome and lead to technical artifacts in downstream analyses and applications. Heteromorphic sex chromosomes, in particular, present a case of sequence homology likely to affect all individuals in a given species.

Sex chromosomes in therians—the clade containing eutherian mammals and marsupials—share a common evolutionary origin as a pair of homologous autosomes (Glas *et al*., 1999). Approximately 180 to 210 million years ago, they began differentiating from each other through a series of recombination suppression events and subsequent gene loss on the Y chromosome (Rens *et al*., 2007; Lahn and Page, 1999; Livernois *et al*., 2012; Wilson Sayres and Makova, 2013). However, this pattern is not unique to mammalian evolution or even XX/XY systems, and occurs often across taxa with genetic sex determination (Bergero and Charlesworth, 2009; Wilson and Makova, 2009). This shared origin and complex history characteristic of sex chromosomes lead to unique challenges for genome assembly and analysis, including large blocks of homologous sequence between the sex chromosomes—called gametologous sequence— that we hypothesize can lead to the mismapping of reads between the sex chromosomes. Further, the sex chromosomes of many species contain pseudoautosomal regions (PARs; of which humans have two: PAR1 and PAR2)—regions that have not differentiated between the sex chromosomes and are identical in sequence between the two sex chromosomes (Simmler *et al*., 1985; Ross *et al*., 2005). A reference genome that includes the entire sequence content from both sex chromosomes will thus duplicate the PARs and substantially reduce mapping quality in these regions because most reads will identically map to two regions in the reference assembly. This stands in contrast to autosomal sequence, for which each diploid autosome is represented just once in the reference genome.

The technical challenges presented by the biological realities of the sex chromosomes can lead to erroneous genotype calls, so the sex chromosomes are routinely excluded from genome-wide analyses (e.g., Wise *et al*., 2013). This is unfortunate because the sex chromosomes contribute to phenotype and disease etiology (e.g., Chang *et al*., 2014) and are useful in population genetic inference of demography and patterns of natural selection (Webster and Wilson Sayres, 2016; Wilson Sayres, 2018; Vicoso and Charlesworth, 2006; Ellegren, 2009; Meisel and Connallon, 2013).

A number of tools, methods, and frameworks have been developed to aid in the identification of sex-linked sequence (e.g., Muyle *et al*., 2016), inference of an individual’s sex chromosome complement (e.g., Madel *et al*., 2016), and handling of some of the technical challenges sex chromosomes present in genome-wide association studies (e.g., Gao *et al*., 2015). However, to our knowledge, there is no tool that simultaneously facilitates the identification of sex chromosome complement and corrects for associated technical biases for the purposes of short read mapping and variant calling.

Out of the urgent need to understand the effects of sex chromosome homology on next-generation sequencing analyses, in this paper we first test whether sequence homology between sex chromosomes can confound aspects of read mapping and lead to downstream errors in sequence analysis. We then present XYalign, a tool developed to perform three major tasks: (1) aid in the characterization of an individual’s sex chromosome complement; (2) identify and correct for technical artifacts arising from sex chromosome sequence homology; and (3) tabulate and visualize important metrics for quality control such as mapping quality, sequencing depth, and allele balance. We show how XYalign can be used to identify XX and XY individuals across sequencing depths and capture techniques. We also show that the default steps taken by XYalign correct many mismapped reads on the sex chromosomes, resulting in more accurate variant calling. Finally, because XYalign is designed to be both scalable and customizable, we discuss how it can be used in a variety of situations including genetic sex identification in both XX/XY and ZZ/ZW systems, identification of sex-linked sequences and pseudoautosomal regions in new draft genomes, correction of technical biases in genomic and transcriptomic data, detection of aneuploidy, and investigation of mapping success across arbitrary chromosomes.

## Methods

### Implementation

XYalign is implemented in Python and uses a number of third-party Python packages including Matplotlib (Hunter, 2007), NumPy (Oliphant, 2006), Pandas (McKinney, 2010), PyBedTools (Quinlan and Hall, 2010; Dale *et al*., 2011), PySam (https://github.com/pysam-developers/pysam), and SciPy (Jones *et al*., 2001). It further wraps the following external tools: repair.sh and shuffle.sh from BBTools (https://sourceforge.net/projects/bbmap/), BWA (Li, 2013), Platypus (Rimmer *et al*., 2014), Sambamba (Tarasov *et al*., 2015), and SAMtools (Li *et al*., 2009).

XYalign is composed of six modules that can be called individually or serve as steps in a full pipeline: PREPARE_REFERENCE, CHROM_STATS, ANALYZE_BAM, CHARACTERIZE_SEX_CHROMS, STRIP_READS, and REMAPPING. Below, we discuss each module as a step in the full XYalign pipeline using human samples (XX/XY sex determination) as an example. Note, however, that XYalign will work with other sex chromosome systems (e.g., ZZ/ZW) and on arbitrary chromosomes (e.g., detecting autosomal aneuploidy). We describe examples of XYalign commands in the section titled “Use Cases.”

The PREPARE_REFERENCE module generates two versions of the same reference genome: one for the homogametic sex (e.g., XX) and one for the heterogametic sex (e.g, XY). In the simplest case, it will completely hard-mask the Y chromosome with Ns in the XX version of the reference. Optionally, it will also accept one or more BED files containing regions to hard mask in both reference versions. If pseudoautosomal regions (PARs) are present on both sex chromosome sequences in the reference, we strongly suggest masking the PARs on the Y chromosome, allowing reads from these regions to map exclusively to the X chromosome in XY individuals. In XYalign, we use hard masks, rather than omitting the Y chromosome in the XX reference version because these hard masks allow files from both references to share the same sequence dictionaries and indices, thus permitting seamless integration of files from both references into downstream analyses (e.g., joint variant calling).

The CHROM_STATS module provides a relatively quick comparison of mapping quality and sequencing depth across one or more chromosomes and over multiple BAM files. While this provides a less detailed perspective than ANALYZE_BAM or CHARACTERIZE_SEX_CHROMS (detailed below), we envision it to be especially useful in at least two different cases. First, in well-characterized systems (e.g., human), comparing chromosome-wide values of mean mapping quality and depth represent a quick and often sufficient way to identify the sex chromosome complement (e.g., XX or XY) of individuals across a population. Second, in uncharacterized systems, the CHROM_STATS output provides information that can help with the identification of sex-linked scaffolds. It is important to note, however, that results for both cases will vary based on ploidy and with differences in the degree of sequence homology between the sex chromosomes.

The ANALYZE_BAM module runs a series of analyses designed to aid in the identification of sex-linked sequence and characterize the sex chromosome content of an individual. In doing so, it provides more detailed metrics than CHROM_STATS. For ANALYZE_BAM, XYalign runs Platypus (Rimmer *et al*., 2014) across multiple threads, if permitted, to identify variants. It then parses the output VCF file containing the variants, applies filters for site quality, genotype quality, and read depth, and plots the read balance at variant sites. Here, we define read balance at a given site as the number of reads containing the alternate allele (i.e., nonreference allele) divided by the total number of reads mapped to the position. XYalign produces plots and tables for read balance per site, as well as mean read balance and variant count per genomic bin or window across a chromosome. We anticipate these data will not only be useful for masking regions containing incorrect genotypes but will also aid in the identification of PARs as well. XYalign next traverses the BAM file, calculating mean mapping quality and an approximation of mean depth in windows across the genome. During traversal, secondary and supplementary read mappings are ignored, and depth is calculated as the total length of all reads mapping to a genomic window divided by the total length of the window. We have found that this heuristic approximation is very similar to calculations of exact depth, particularly as window sizes increase, and is much faster to compute across entire chromosomes. XYalign will output a table containing genomic coordinates, mean depth, and mean mapping quality for each window. It will then filter windows based on user-defined thresholds of mean depth and mapping quality and output two BED files containing windows that passed and failed these thresholds, respectively, which can be used for additional masking in downstream applications. Finally, XYalign will output plots of mapping quality and depth in each window across each chromosome.

After running ANALYZE_BAM, the windows meeting thresholds can be used by the CHARACTERIZE_SEX_CHROMS module to systematically compare mean depth in pairs of chromosomes using three different approaches. The first is a bootstrap analysis that provides 95% confidence intervals of mean window depth for each of the chromosomes in a given pair to test for overlap. The second is a permutation analysis to test for differences in depth between the two chromosomes. The third is a two-sample Kolmogorov-Smirnov test (Massey Jr., 1951). Though all three tests are implemented in XYalign, we only present results from the bootstrap analyses in this manuscript. Further, while we present analyses pairing sex chromosomes with an autosome (here we use chromosome 19), the chromosome pairs are arbitrary and can feature any scaffolds or chromosomes in a reference genome, depending on a user’s needs.

Finally, the REMAPPING module will infer the presence or absence of a Y chromosome based on the results of CHARACTERIZE_SEX_CHROMS. If a Y chromosome is not detected, the STRIP_READS module will iteratively remove reads from the sex chromosomes by read group ID using SAMtools (Li *et al*., 2009), writing FASTQ files for each. XYalign will use repair.sh from BBTools to sort and re-pair paired-end reads or shuffle.sh from BBTools to sort single-end reads for each read group. The REMAPPING module then maps reads with BWA-MEM (Li, 2013) and sorts alignments with SAMtools (Li *et al*., 2009) by read group. If more than one read group is present, the resulting BAM files are merged using SAMtools (Li *et al*., 2009). Finally, XYalign uses Sambamba (Tarasov *et al*., 2015) to isolate all scaffolds not associated with sex chromosomes from the original BAM file and then SAMtools (Li *et al*., 2009) to merge this file with the BAM file containing the new sex chromosome mappings.

When run as a full pipeline on a sample, XYalign will first call PREPARE_REFERENCE to generate XX and XY reference genomes with appropriate masks. Next, it will call ANALYZE_BAM and CHARACTERIZE_SEX_CHROMS to preliminarily analyze the unprocessed input BAM file. Then, based on the results of CHARACTERIZE_SEX_CHROMS, XYalign will call STRIP_READS to extract reads from the sex chromosomes and REMAPPING to remap to the appropriate reference genome output from PREPARE_REFERENCE. Finally, XYalign will re-run the ANALYZE_BAM module to analyze the remapped BAM file and provide metrics to allow a before-and-after comparison.

While we anticipate that this full pipeline will be useful in certain situations, it is neither the only nor the best-suited option for most users. Rather, we expect that most users will call modules individually. We give examples of other implementations below and provide recommendations for incorporating XYalign into bioinformatic pipelines in the discussion.

### Operation

XYalign is available via PyPI (https://pypi.python.org/pypi), Bioconda (Grüning *et al*., 2017), and Github (https://github.com/WilsonSayresLab/XYalign), with documentation hosted at Read the Docs (http://xyalign.readthedocs.io/en/latest/). A full environment containing all dependencies can be most easily installed and managed using Anaconda (https://www.continuum.io/) and Bioconda (Grüning *et al*., 2017). It has been tested on a variety of UNIX operating systems (including Linux and MacOS), but it is not currently supported for the Windows operating system.

XYalign is typically invoked from the command line, but it can be imported into Python scripts for more customized use cases. The next section lists a number of example commands that illustrate how to call the full pipeline as well as individual modules.

### Use Cases

To highlight some features of XYalign and its flexibility, we used two datasets from publicly available sources (Supplemental Table S1): (1) exome, low-coverage whole-genome, and high-coverage whole-genome sequencing data from one male (HG00512) and one female (HG00513) from the 1000 Genomes Project (Dataset 1; (The 1000 Genomes Project Consortium, 2015); and (2) 24 high-coverage whole genomes from the 1000 Genomes Project (Dataset 2; (Sudmant *et al*., 2015). For Dataset 1, we mapped reads to the hg19 version of the human reference genome (International Human Genome Sequencing Consortium, 2001) using BWA MEM (Li, 2013), marked duplicates with SAMBLASTER (Faust and Hall, 2014), and used SAMtools (Li *et al*., 2009) to sort, index, and merge BAM files. The publicly available BAM files for Dataset 2 were previously mapped using a different version of hg19 (from the Broad Institute’s GATK Resource Bundle; https://software.broadinstitute.org/gatk/download/bundle), which we used for analyses involving this dataset.

With these datasets, we examined three potential uses of XYalign. First, to explore the effects of simple corrections for technical biases arising from sequence homology on the sex chromosomes, we ran the full XYalign pipleline on all six BAM files from Dataset 1. In all cases below, the exact commands are included in the Supplementary Material and in a Snakemake (Köster and Rahmann, 2012) pipeline available with the XYalign software distribution (Webster *et al*., 2018), and templates are shown here for convenience. Because we were using the same output directory for these analyses, we avoided conflicts by initially preparing separate XX and XY references using the following command:

> *xyalign --PREPARE_REFERENCE --ref <hg19 reference genome> --xx_ref_out hg19.XXonly.fasta --xy_ref_out hg19.XY.fasta --output_dir <output_directory> -- x_chromosome chrX --y_chromosome chrY --bwa_index True*

where *<hg19 reference genome>* was the path to the FASTA file containing the hg19 reference, *<input bam file>* was a sorted BAM file, and *<output directory>* was the directory where XYalign wrote output. We then ran the full pipeline on all six files using the following command template:

> *xyalign --ref <hg19 reference genome> --bam <input bam file> --output_dir <output directory> --sample_id <sample ID> --cpus 4 --reference_mask hg19_PAR_Ymask_startEnd. bed --window_size 5000 --chromosomes chr19 chrX chrY -- x_chromosome chrX --y_chromosome chrY --xmx 4g --fastq_compression 4 -- min_depth_filter 0.2 --max_depth_filter 2 --xx_ref_in hg19.XXonly.fasta --xy_ref_in ref_out hg19.XY.fasta*,

where *<sample ID>* was the identification code for a given sample, *hg19_PAR_Ymask_startEnd.bed* was a BED file containing the genomic coordinates of the PARs in the hg19 assembly, and *hg19.XXonly.fasta* and *hg19.XY.fasta* were the two FASTA formatted reference genomes prepared in the previous step.

Next, we examined how the metrics generated by XYalign can be used to identify the sex chromosome complement of individuals with both datasets. Here, we used the CHARACTERIZE_SEX_CHROMS module of XYalign. This was automatically done for Dataset 1 when running the full pipeline (see above). For Dataset 2, we used the following command template for BAM files:

> *xyalign --CHARACTERIZE_SEX CHROMS --ref <1000 genomes reference genome> --bam <input bam file> --output_dir <output directory> --sample_id <sample ID> --cpus 4 --window_size 5000 --chromosomes 19 X Y --x_chromosome X --y_chromosome Y*

Finally, we explored the utility of the CHROM_STATS module for identifying sex chromosome complement and potentially sex-linked scaffolds with both datasets using the following command template for BAM files:

> *xyalign --CHROM_STATS --chromosomes chr1 chr8 chr19 chrX chrY chrM --bam <input_bam_1> <input_bam_2> <input_bam_3> --ref null --sample id <name_of_analysis> --output dir <output_dir>*

We additionally ran CHROM_STATS using the above command with the addition of the “--use_counts” flag to calculate metrics using only the number of reads mapping to each chromosome.

We visualized all CHROM_STATS results using the plot_count_stats utility, with the command template:

> *plot_count_stats --input <chrom_stats output file> --output_prefix <output prefix>-- meta <metadata text file< --exclude_suffix <suffix> --first_chr chrX --second_chr chrY --const_chr chr19 --var1 marker color --var1 marker_vals darklateblue thistle -- var2_marker shape --var2_marker_vals o s v --marker_size 1700 --legend_marker_scale 0.4*

where *<chrom_stats_output_file>* was either the count, mapping quality, or depth output of CHROM_STATS, *<metadata text file>* was the appropriate metadata text file, and *<suffix>* was the string to remove from filenames.

### Sex chromosome coordinates

To better understand the genomic context of technical artifacts, we explored variation in mapping quality and depth in association with genomic features on the X and Y chromosomes. On the Y chromosome, we used coordinates from Poznik *et al*. (2013) based on Skaletsky *et al*. (2003) (provided by D. Poznik, personal communication). On the X chromosome, we obtained coordinates for ampliconic regions from Cotter *et al*. (2016) and all other regions (PARs, telomeres, centromere, and XTR) from the UCSC Table Browser (Karolchik *et al*., 2004). We define the X-transposed region (XTR) on the X chromosome as beginning at the start of DXS1217 and ending at the end of DXS3 (Mumm *et al*., 1997).

To count variants falling in major genomic regions, we intersected a BED file containing coordinates with VCF files using BEDTools (Quinlan and Hall, 2010). We first filtered VCF files using BCFtools (Li *et al*., 2009) with the following command template:

> *bcftools filter --include ‘INFO/MQ>=30 && %QUAL>=30’ <input_vcf>*

We then identified variants unique to each file through iterations of the “subtract” command in BEDtools (Quinlan and Hall, 2010):

> *bedtools subtract -header -a <first_vcf> -b <second_vcf>*

Finally, in each region, we counted variants present in a given filtered VCF file using the BEDtools (Quinlan and Hall, 2010) “intersect” command:

> *bedtools intersect -c -a <BED file> -b <vcf_file>*

where *<BED_file>* is the BED file containing genomic coordinates (Supplemental Table S2).

### Specific commands

We provide Snakemake (Köster and Rahmann, 2012) workflows for all assembly and analysis steps on Github (https://github.com/WilsonSayresLab/XYalign), Zenodo (Webster *et al*., 2018), and in the Supplemental Material.

## Results and Discussion

We observed a number of artifacts stemming from several methodological challenges presented by the human sex chromosomes. First, PAR1 and PAR2 on both sex chromosomes are clearly identifiable in genomic scatter plots of mapping quality and depth in all datasets (Figures 1-3). While these results are not surprising given the sequence homology in these regions (Ross *et al*., 2005), they highlight the fact that these measures can help identify other similarly problematic areas. For example, there is a region of reduced mapping quality on the X chromosome beginning near 88.4 Mb and ending near 92.3 Mb (Figure 2). This corresponds to the X-transposed region (XTR), which arose by a duplication from the X to the Y chromosome in the human lineage since its divergence with the chimpanzee-bonobo lineage (Page *et al*., 1984; Ross *et al*., 2005). This region retains more than 98% sequence similarity between the X and Y chromosome (Ross *et al*., 2005), likely leading to the reduction in mapping quality. Interestingly, we observe a similar decrease in mapping quality on the Y chromosome beginning near 2.9 Mb and ending near 6.6 Mb, corresponding with known coordinates of the XTR on the Y chromosome (Figure 3). In fact, integrating mapping quality and depth recapitulates established genomic features of both sex chromosomes (e.g., ampliconic regions, PARs, and XTRs) described in previous studies (Figures 1-3; Poznik *et al*., 2013; Mueller *et al*., 2013). This suggests that, in at least some cases, the output of XYalign can be used to quickly explore broad patterns of genomic architecture and mask regions likely to introduce technical difficulties in genomic analyses.

**Figure 1.**
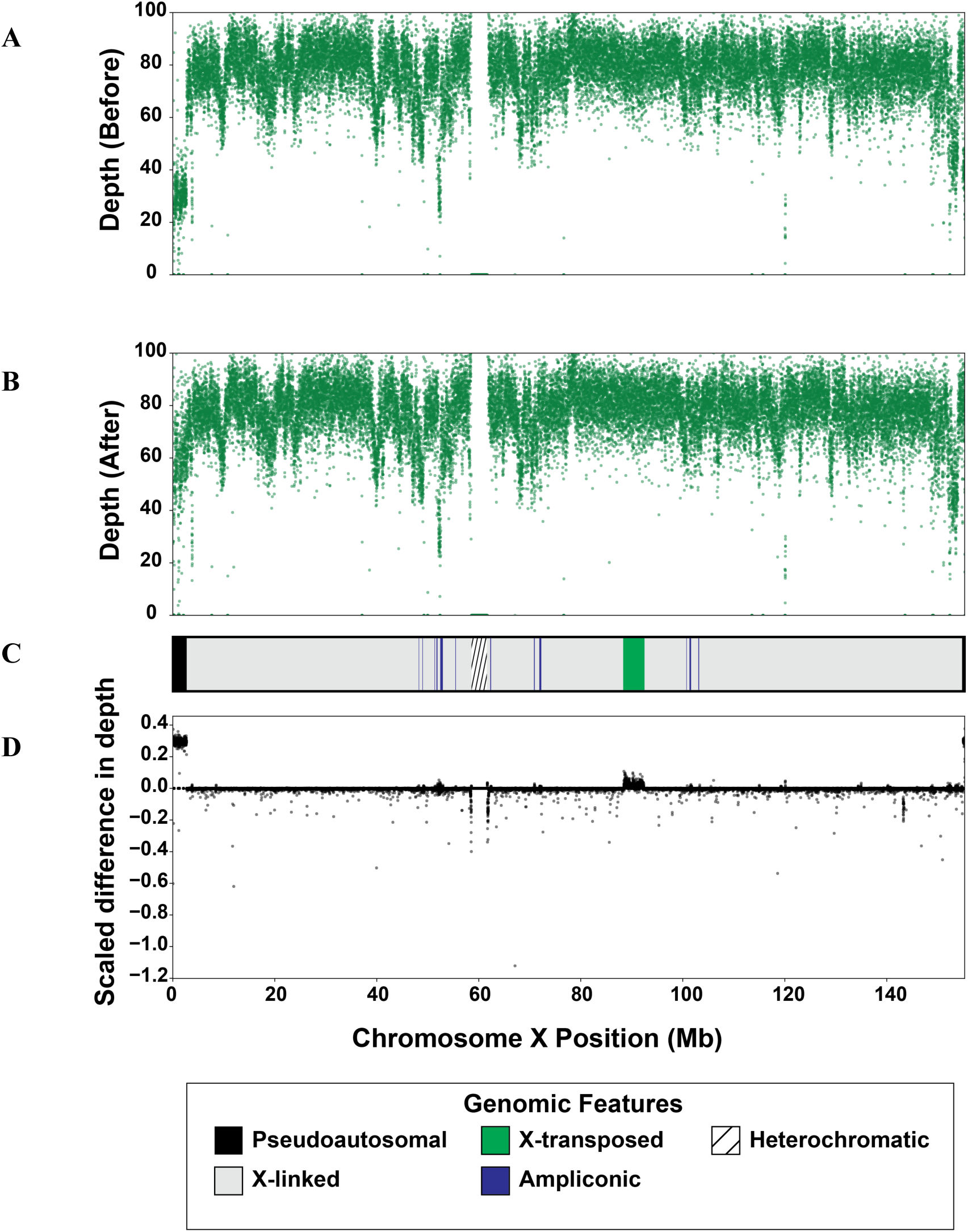
Sequencing depth on chromosome X before and after XYalign. Mean sequencing depth in 5 kb windows across the X chromosome before (A) and after (B) XYalign processing. Changes in depth (D) are presented as the sign of the difference times the absolute value of the log_10_ difference, where the difference is depth after XYalign minus depth before XYalign. The chromosome map (C) presents the location of X chromosome genomic features depicted in the legend. X chromosome coordinates are identical in all plots.

**Figure 2.**
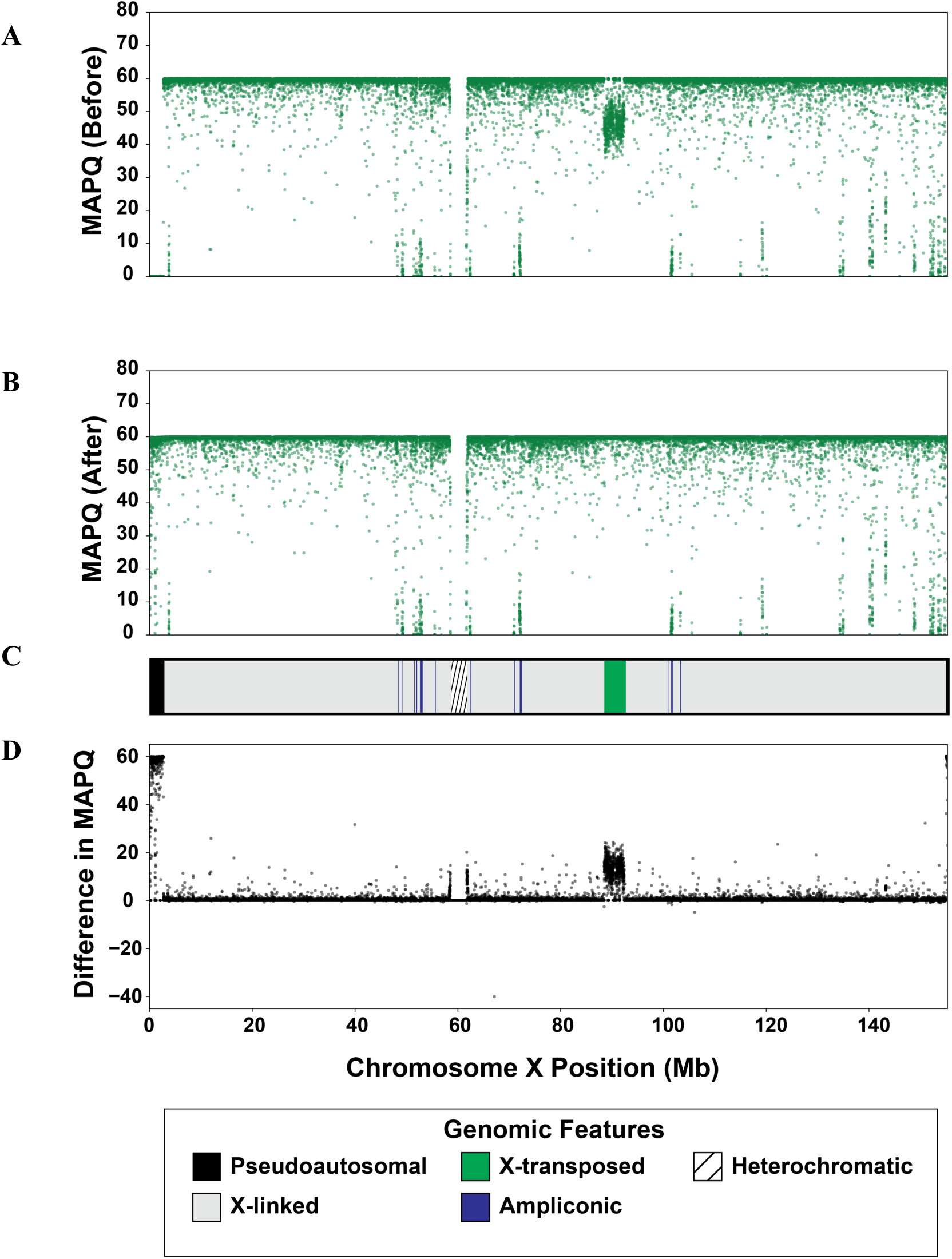
Mapping quality on chromosome X before and after XYalign. Mean mapping quality (MAPQ) in 5 kb windows across the X chromosome before (A) and after (B) XYalign processing. Changes in MAPQ (D) are presented as the difference is MAPQ after XYalign minus MAPQ before XYalign. The chromosome map (C) presents the location of X chromosome genomic features depicted in the legend. X chromosome coordinates are identical in all plots.

**Figure 3.**
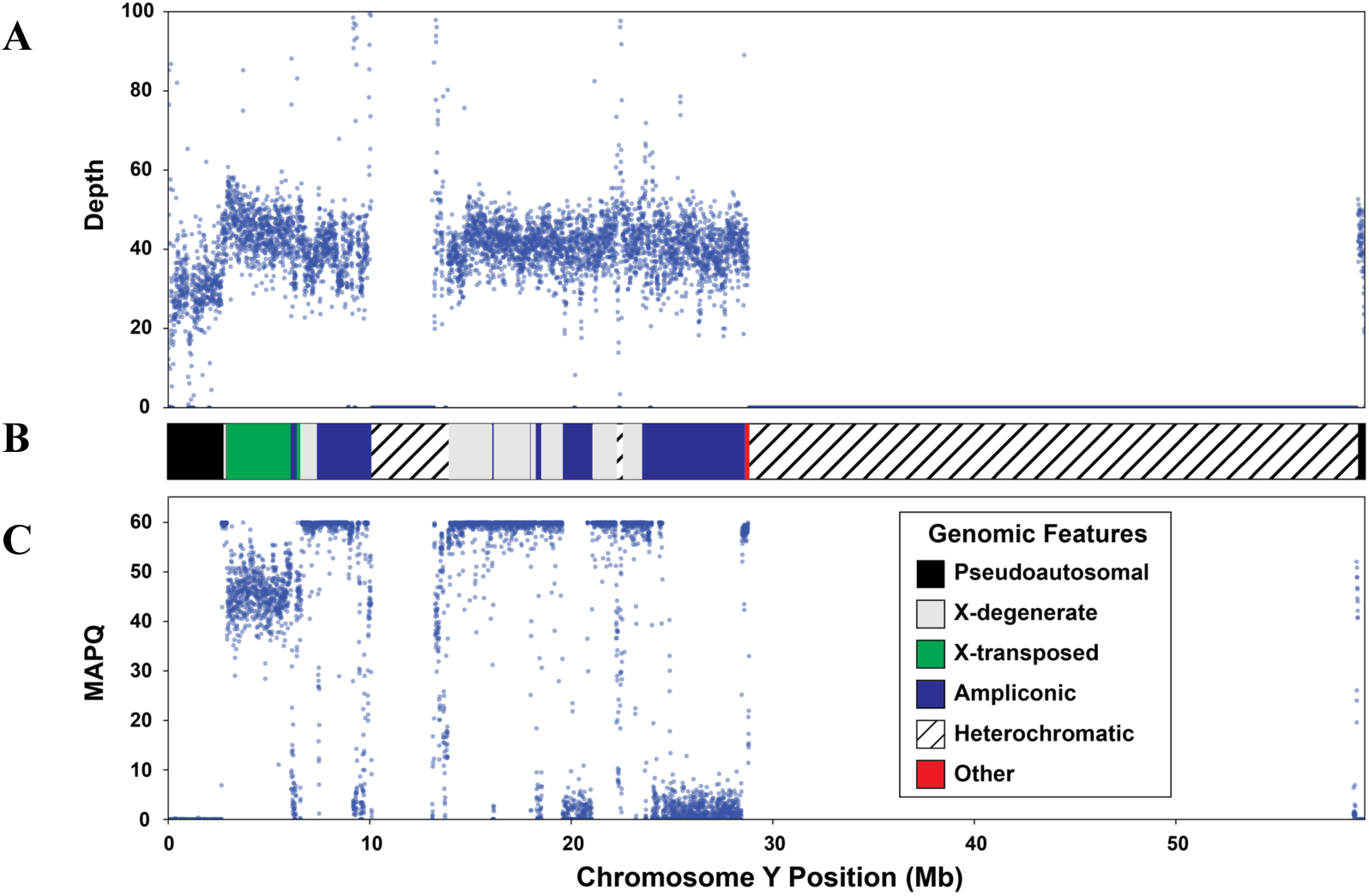
Y chromosome sequencing depth and quality. Mean sequencing depth (A) and mapping quality (MAPQ; C) in 5 kb windows across the Y chromosome. The chromosome map (B) presents the location of Y chromosome genomic features depicted in the legend. Y chromosome coordinates are identical in all plots.

By hard-masking the Y chromosome in the XX reference genome, and the pseudoautosomal regions (PAR1 and PAR2) in the reference genome for the XY reference genome, we observed clear improvements in read mapping (Figures 1-2). On the X chromosome, all metrics exhibited striking improvements in PAR1, PAR2, and XTR (Figures 1 and 2). Furthermore, the Y chromosome of the XX individual no longer exhibited any variant calls or mapped reads, though many passed filters before processing (variants before: 4266; variants after: 0; mapped reads before: 5,729,007; reads mapped after: 0). While this is expected given the hard masking of the Y chromosome, it is worth emphasizing that this is consistent with the biological state of the individual.

We found that these improvements in mapping on the X chromosome after masking the Y chromosome substantially impacted downstream variant calling (Table 1). Unsurprisingly, the effect was most pronounced in the PARs, in which thousands of variants were callable after masking the identical sequences present on the Y chromosome in the reference assembly. The XTR also had a large increase in the number of variants detected after Y masking—an average of 85.4 variants per megabase of sequence (Table 1). However, effects were not limited to these regions of well-documented homology: both the X-added region (XAR) and X-conserved region (XCR) contained hundreds of affected variants, suggesting effects of more extensive homology across the sex chromosomes.

**Table 1.** The effect of sex chromosome homology on variant calling on the X chromosome.

| Region<sup>a</sup> | Length<sup>b</sup> | False positives (per Mb)<sup>c</sup> | False negatives (per Mb)<sup>d</sup> |
| --- | --- | --- | --- |
| PAR1 | 2,589,520 | 0 (0) | 7563 (2920.6) |
| PAR2 | 329,516 | 0 (0) | 633 (1921) |
| XTR | 4,287,237 | 40 (9.3) | 366 (85.4) |
| XAR | 55,982,492 | 299 (5.3) | 400 (7.2) |
| XCR | 89,011,795 | 610 (6.9) | 523 (5.9) |
| *Total* | *152,250,560* | *949 (6.2)* | *9485 (62.3)* |

<sup>a</sup>PAR1: pseudoautosomal region 1; PAR2: pseudoautosomal region 2; XTR: X-transposed region; XAR: X-added region; XCR: X-conserved region.
<sup>b</sup>Total sequence length of region in base pairs.
<sup>c</sup>Total number of false positive variants after filtering, defined as being present before but not after Y chromosome masking. Variants per Mb of sequence are presented in parentheses.
<sup>d</sup>Total number of false negative variants after filtering, defined as being present after but not before Y chromosome masking. Variants per Mb of sequence are presented in parentheses.

### Inferring Genetic Sex

In our analyses, the most striking measure for assessing an individual’s sex chromosome complement was the distribution of read balances across a chromosome (Figure 4). Specifically, when we plotted the distribution of the fraction of reads containing a nonreference allele at a given variant site, we observed that diploid chromosomes (e.g., autosomes, and chromosome X in XX individuals) exhibited peaks both around 0.5 and 1.0, consistent with the presence of heterozygous sites and sites homozygous for a nonreference allele, respectively (Figure 4). In the case of the X chromosome in XY individuals, we observed a single peak near 1.0, consistent with an expected haploid state (i.e., no heterozygous sites; Figure 4). We observed one exception to this pattern: the Y chromosome exhibited a peak around 0.2 in addition to the one near 1.0 (Figure 4). All variants included in analyses met thresholds for depth, site quality, and genotype quality, so quality does not appear to be a driving factor of this pattern. This pattern also remained after genomic windows of low mapping quality and irregular depth were removed. We are currently unable to explain these results and more work is thus required to understand the factors responsible for this pattern and whether similar results are obtained on the W chromosome in ZW systems.

**Figure 4.**
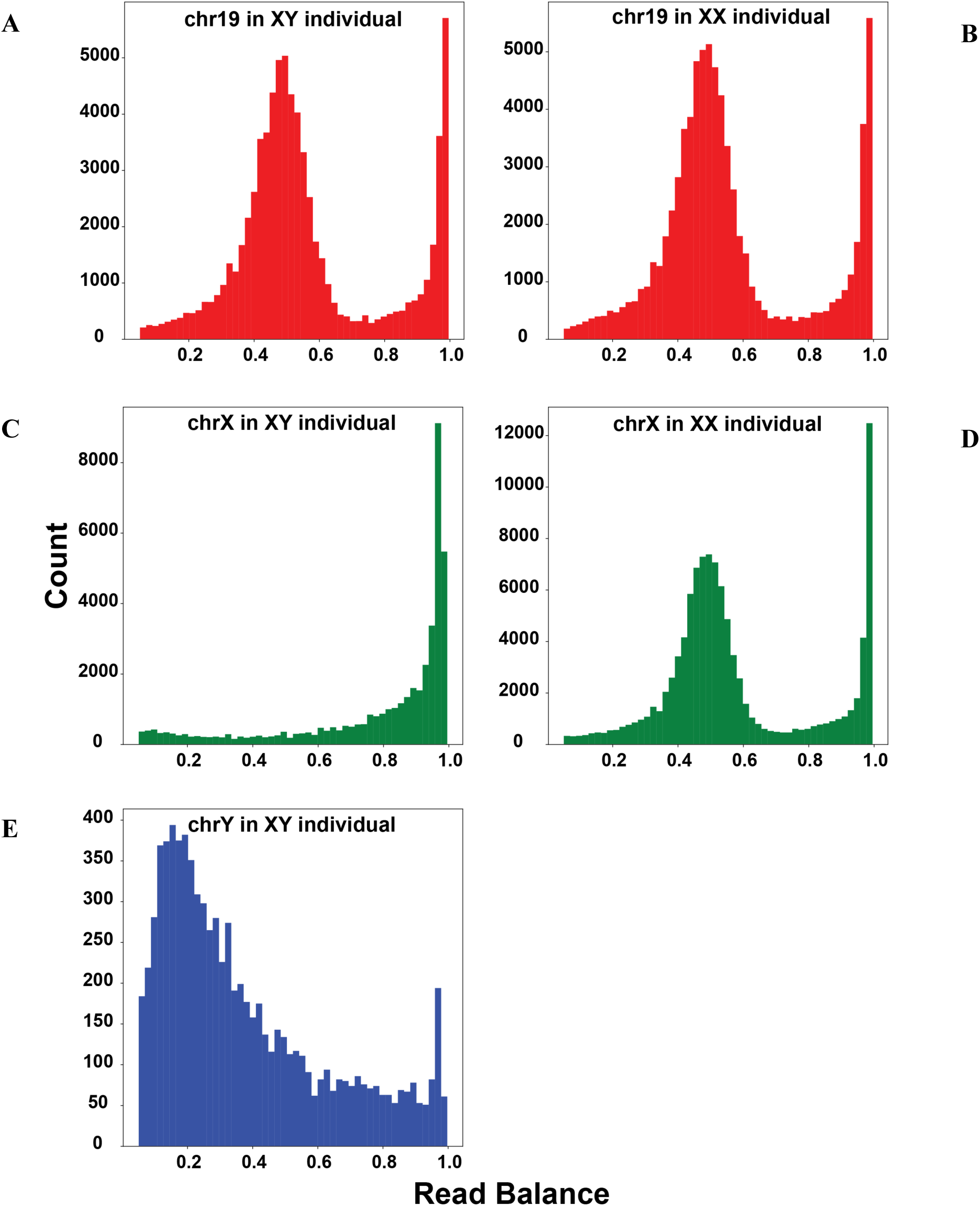
Read balance in XY and XX samples. Histograms of read balance for an XY sample (Left Column; A, C, and E) and XX sample (Right Column; B and D) across chromosome 19 (Top; A and B), chromosome X (Middle; C and D), and chromosome Y (Bottom; E). Read balance at a given site is defined as the number of reads containing a non-reference allele divided by the total number of reads mapped to a site. Read balances between 0.05 and 1.0 are presented.

Across datasets, we observed variation in relative depth of the X and Y chromosomes in XX and XY individuals, particularly among different sequencing strategies: exome, low-coverage whole-genome, and high-coverage whole-genome sequencing (Figure 5A). However, within datasets, XX and XY individuals were clearly differentiated (Figure 5; Supplemental Figure S1). This pattern suggests that a general threshold for assigning different genetic sexes across a range of organisms and sequencing experiments might be difficult to implement. That being said, within species, some combination of depth, mapping quality, and read balance is likely to be informative. For example, in humans, relative mapping quality appears to be informative in some sequencing strategies, particularly exome sequencing (Figure 5B). This should be explored in each experiment, however, as we did not observe this differentiation in the uncorrected 1000 Genomes high-coverage samples (Supplemental Figure S2).

**Figure 5.**
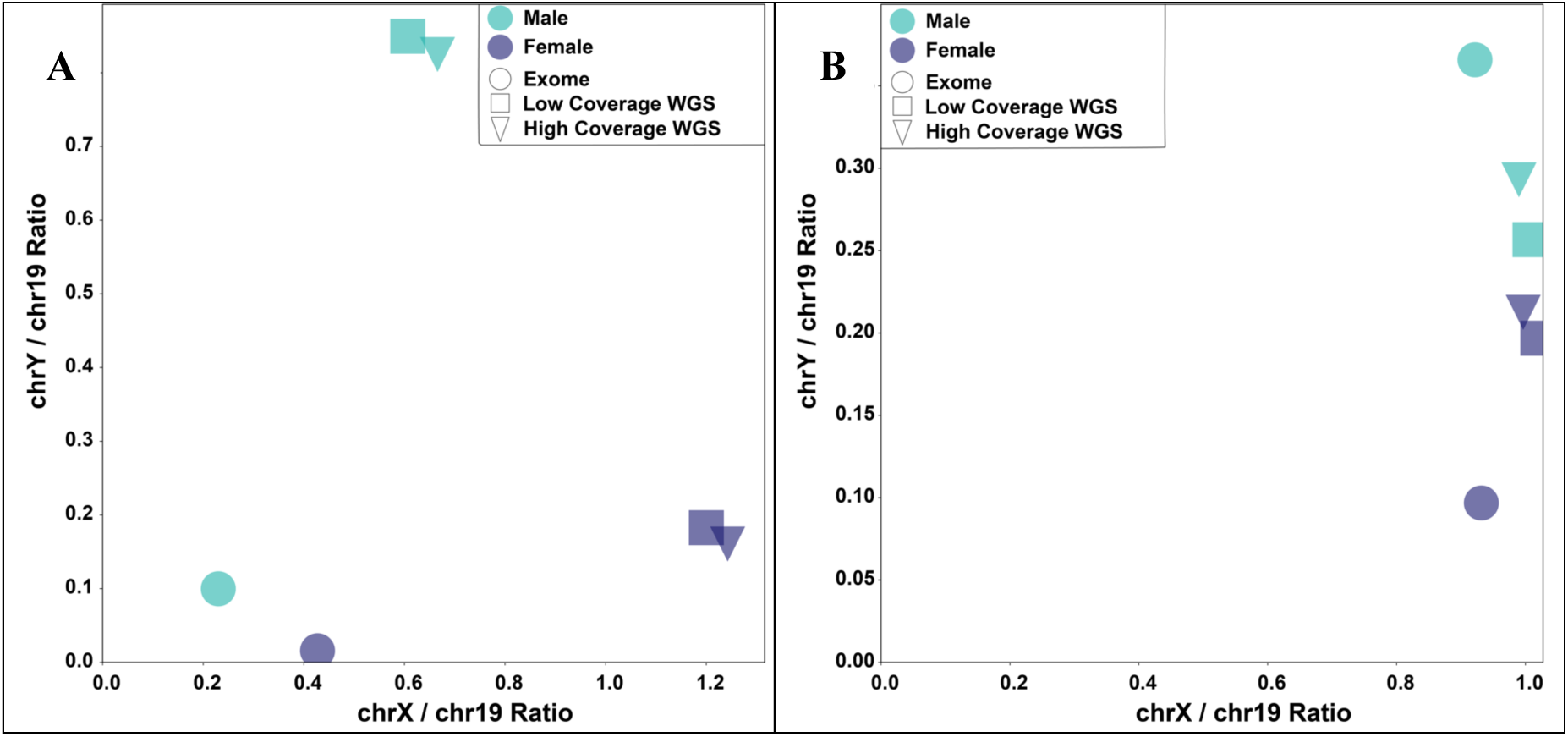
Relative sequencing depth and mapping quality on the X and Y chromosomes across different sequencing strategies. Values of relative (A) sequencing depth and (B) mapping quality come from exome (circles), low-coverage whole-genome sequencing (squares), and high-coverage whole-genome sequencing (triangles) for a single male (green) and female (blue) individual. Mean depth and MAPQ on chromosome 19 was used to normalize the sex chromosomes.

Generating these results for all individuals in a study is easy to do with XYalign: one can iteratively run the CHARACTERIZE_SEX_CHROMS module on preliminarily mapped BAM files. Then, the results from all individuals can be analyzed together (see the Supplementary Material for an example of such analysis). At least with human samples, for which X and Y chromosomes are very differentiated, this process can be sped up significantly with the CHROM_STATS module. In our data, read counts on the X and Y chromosomes quickly and clearly clustered male and female samples within sequencing strategies (i.e., exome, low-coverage whole-genome, and high-coverage whole-genome; Supplemental Figures S3-S4). However, the success of this procedure likely depends on the degree of differentiation between sex chromosomes; other organisms might require the statistics output as part of the CHARACTERIZE_SEX_CHROMS module.

### Recommendations for researchers

Based on these results, we can make the following recommendations for researchers. For organisms with multiple sex chromosomes assembled (e.g., both X and Y or both Z and W) and included in reference assemblies (e.g., human, chimpanzee, rhesus macaque, gorilla, mouse, rat, chicken, *Drosophila), if the genetic sex of every individual is known*, the user may: (1) prepare separate assemblies for the different sexes using the PREPARE_REFERENCE module; (2) map and process reads according to user’s typical pipeline (mapping individuals by sex to their corresponding reference); (3) confirm genetic sex using the CHROM_STATS module; (4) remap any incorrectly assigned individuals; and (5) proceed with downstream analyses. *If genetic sexes of individuals are unknown*, the user should then: (1) prepare separate assemblies for the different sexes using the PREPARE_REFERENCE module; (2) map and process a suitable number of reads (e.g., whole dataset for exome or a single lane of WGS) according to user’s typical pipeline using the reference genome of the heterogametic sex (i.e., XY or ZW); (3) infer the sex chromosome complement using either CHROM_STATS (for well-characterized and highly divergent sex chromosomes), CHARACTERIZE_SEX_CHROMS, or both; (4) map and process all reads using the prepared reference genome corresponding to the inferred sex of each individual; and (5) run downstream analyses.

For individuals of the homogametic sex (i.e., XX or ZZ), the above recommendations will likely completely remove artifacts stemming from sex chromosome homology, assuming only a single unmasked sex chromosome is left after XYalign processing. However, homology is unavoidable for individuals of the heterogametic sex (i.e., XY or ZW) because both sex chromosomes are required in the reference assembly for mapping. In this case, a more local masking or filtering approach is likely the most promising option. For studies investigating specific variants, for which false negatives are preferable to false positives, we suggest strict variant filtering that includes high thresholds for mapping quality (e.g., thresholds of 55 or higher are required to eliminate the effects of homology in the X-transposed region). However, for studies investigating invariant sites as well (e.g., measures of genetic diversity require information from all monomorphic and polymorphic sites), we recommend filtering entire regions based on, at the very least, mapping and depth metrics. These masks are output by the BAM_ANALYSIS module in XYalign, and for this use, we recommend using small windows (e.g, 1 kb to 5 kb) and exploring a variety of depths. Finally, in all cases, if pseudoautosomal regions are present in the reference genome, they should be masked in the heterogametic sex’s assembly output by the PREPARE_REFERENCE module.

### Additional uses for XYalign

While the development of XYalign was motivated by challenges surrounding erroneous read mapping and variant calling due to sex chromosome homology in human sequencing experiments, the software can be utilized in a number of additional scenarios. First, it can be applied to any species with heteromorphic sex chromosomes to identify relative quality and depth. The results output by CHROM_STATS, ANALYZE_BAM, and CHARACTERIZE_SEX_CHROMS can be used to identify sex-linked scaffolds, characterize sex chromosome complements, and determine the most appropriate remapping strategy. Second, XYalign can be used to detect relative sequencing depth, mapping quality, and read balance on any chromosome, not just the sex chromosomes. In addition to exploring mapping artifacts, we anticipate that this will aid in detection of aneuploidy in the autosomes. However, we note that many programs exist to calculate depth of coverage (e.g., Quinlan and Hall, 2010; Pedersen and Quinlan, 2018; McKenna *et al*., 2010) and identify structural variants within statistical frameworks (e.g., Chen *et al*., 2016; Layer *et al*., 2014; Abyzov *et al*., 2011; Roller *et al*., 2016). Accordingly, XYalign might not be the most appropriate option for detecting local phenomena such as copy number variants. Finally, XYalign may also be extended to other types of data, including RNA sequencing data, where the same fundamental challenge (gametologous sequence between the X and Y) can affect mapping and variant calling. In particular, we expect biases to manifest in differential expression and biased-allelic expression, and suggest that the PREPARE_REFERENCE module be considered for all RNA sequencing experiments in systems with sex chromosomes.

## Conclusion

We showed that the complex evolutionary history of the sex chromosomes creates mapping biases in next-generation sequencing data that have downstream effects on variant calling and other analyses. These technical artifacts are likely present in most genomic datasets of species with chromosomal sex determination. However, many of these biases can be corrected through the strategic use of masks during read mapping and the filtering of variants. We developed XYalign, a tool that facilitates the characterization of an individual’s sex chromosome complement and implements this masking strategy to correct these technical biases. We illustrated how XYalign can be used to identify the presence or absence of a Y chromosome, characterize mapping biases across the genome, and correct for these mapping biases. XYalign provides a framework to generate more robust short read mapping and improve variant calling on the sex chromosomes.

### Software Availability

XYalign is available on Github (https://github.com/WilsonSayresLab/XYalign) under a GNU General Public License (version 3). We have also deposited a static version of the source code used for analyses in this paper at Zenodo (Webster *et al*., 2018).

## Author Contributions

MAWS and THW conceived the research. All authors participated in the initial design of the software. THW was responsible for subsequent design, development, and implementation of the software. BG, EK, TNP, WW, and THW tested the software. THW analyzed the data. THW and MAWS wrote the manuscript. All authors were involved in the revision of the manuscript and have agreed to the final content.

## Acknowledgements

We thank the organizers of Hackseq 2016 for facilitating this project and supporting this collaboration; members of the Wilson Sayres lab for helpful comments; and ASU Research Computing for computational resources.

